# Controlled tumor heterogeneity in a co-culture system by 3D bio-printed tumor-on-chip model

**DOI:** 10.1101/2023.05.02.539106

**Authors:** Nafiseh Moghimi, Seied Ali Hosseini, Altay Burak Dalan, Dorsa Mohammadizadeh, Aaron Goldman, Mohammad Kohandel

**Affiliations:** Department of Applied Mathematics, University of Waterloo, Waterloo, Canada; Electrical Engineering Department, University of Waterloo, Waterloo, Canada; Department of Medical Genetics, School of Medicine, Yeditepe University, Istanbul, Turkey; Department of Medicine, Harvard Medical School, Boston, MA, USA; Division of Engineering in Medicine, Brigham and Women’s Hospital, Boston, MA, USA

## Abstract

**Background:** Cancer treatment resistance is a consequence of cell diversity and tumor heterogeneity. Tumor cell-cell and cell-microenvironment interactions significantly influence tumor progression and invasion, which have important implications for diagnosis, therapeutic treatment and chemoresistance.

**Method:** In this study, we develop 3D bioprinted in vitro models of the breast cancer tumor microenvironment (TME) made of co-cultured cells distributed in a hydrogel matrix with controlled architecture to model tumor heterogeneity. We hypothesize that the tumor could be represented by a cancer cell-laden co-culture hydrogel construct, whereas its microenvironment can be modeled in a microfluidic chip capable of producing a chemical gradient. Breast cancer cells (MCF7 and MDA-MB-231) and non-tumorigenic mammary epithelial cells (MCF10) were embedded in the alginate-gelatine hydrogels and printed using a multi-cartridge extrusion bioprinter.

**Results:** Our method gives special control on the cell positions in the co-culture system, whereas different tumor architectures can be designed. Cellularly heterogeneous samples comprised of two different cancer cells with controlled density are developed in specific initial locations, i.e. two cell types randomly mixed or positioned in sequential layers. A migration-inducing chemical microenvironment was created in a chamber with a gradual chemical gradient to study the cell migration in the complex tumor construct toward the chemoattractant. As a proof of concept, the different migration pattern of MC7 cells toward the epithelial growth factor gradient was studied with presence of MCF10 in different ratio in this device.

**Conclusion:** Combining 3D bioprinting with microfluidic device in our method provides a great tool to create different tumor architectures as can be seen in different patients, and study cancer cells behaviour with accurate special and temporal resolution.

## Introduction

Breast cancer is the most common cancer in women and the second most common cancer overall.^1^ There were over two million new cases in 2018, and more than 30% of those women died.^1^ The aggressiveness of breast cancer may be due to the known heterogeneity of breast tumors.^2^ Tumor heterogeneity has been observed among patients (inter-tumor heterogeneity) and in each individual tumor (intratumor heterogeneity), which leads to breast cancer aggressiveness and challenges in treatment.^3^ Since tumor may consist of phenotypically different cancer cell populations with different properties, tumor specimen obtained by biopsy does not represent the exact tumor composition. In the case of intratumor heterogeneity, the tumor consists of different phenotypical cell populations, which can be identified by different cell phenotypes, cell density, or their localization in the tumor.^4,5^

Conventional models lack spatial cellular heterogeneity of breast cancer and usually depend on external stimuli or stresses which make them challenging in the formation and study of physiological tumors.^6^ Two-dimensional (2D) culture models also cannot mimic the tumor heterogeneity and microenvironment,^7^ while tumour growth in vivo occurs in a three-dimensional (3D) environment in which the cancer cells are in constant and intimate contact with the extracellular matrix (ECM) and other cells. 3D cancer models development provides economic and ethical benefits for the prediction of tumor response to treatment by reproducing important aspects of tumors, such as the presence of oxygen and nutrients gradients, cell-cell interactions, drug penetration and subpopulations of quiescent cells. Furthermore, 3D in vitro models are bridging the gap between 2D and in vivo studies, thereby reducing the number of animals sacrificed in preclinical studies.^8^ By adding a microfluidic system to a 3D culture in a so-called tumor-on-chip system, recapitulation of the TME gets even more accurate, providing valuable insights into cancer biology. ^9,10^

Bioprinting recently attracted much attention because of its ability to build tissue constructs by precise positioning of cells and biomaterials in a layer-by-layer procedure.^11,12^ In this method, living cells and ECM components are printed together, making a ’bioink’, which can recapitulate the compositional and geometric complexity of the tumor microenvironment (TME) while preserving cell viability.^13–15^ Current bioprinting methods include inkjet, extrusion, and laser-assisted printing.^16,17^ Among them, the extrusion method features versatile material selections and allows tumor heterogeneity to be controlled within the printed matrices by tight spatial control of distinct types of materials at different initial locations.^17^ This highly reproducible process also enables the deposition of materials that have cells of a known density embedded in them. The main advantage of extrusion bioprinting technology is the ability to deposit very high cell densities close to physiological cell densities, which is a major goal of bioprinting methods.^16,17^ Extrusion bioprinting has been successfully used to build human-scale tissue construct,^18^ vascular structures^19,20^ and skin tissues,^21,22^ and results in high printing fidelity and cell viability.

Our goal is to establish a platform for modeling the phenotypic heterogeneity that can occur due to different cell localization in a tumor (tumor center or periphery, uneven oxygen amount) and/or interaction with TME. The present work covers the design and construction of in vitro tumor models via physical and chemical means by the 3D bioprinting of co-cultured cells with controlled distribution in a hydrogel matrix in a microfluidic device. Our hypothesis is that the tumor could be represented by a cancer cell-laden co-culture hydrogel construct, whereas its microenvironment can be modeled in a microfluidic chip capable of producing a chemical gradient. Our composite hydrogel as a bioink comprised of alginate and gelatin, which is compatible to the microscopic architecture of a native tumor stroma. The cell-laden structures which are breast cancer cells embedded in the hydrogels and printed via an extrusion-based bioprinter will create a 3D model tumor that mimics the in vivo microenvironment. We have optimized our bioink based on printability and cell viability measurements. Co-cultured 3D bioprinted constructs are developed with the best chosen bioink. We use a multi-cartridge extrusion bioprinter that allows us to develop cellularly heterogeneous samples comprised of two different breast cancer cells in specific initial locations, specific architecture with controlled density.

Engineering a migration-inducing chemical microenvironment is employed for the creation of an in vitro platform to address metastatic behaviour, which has not yet been fully achieved by current in vitro tumor models. Extracellular chemical gradients are dynamically manipulated via 3D printed constructs containing growth factors in the chamber, enabling post-print cellular modulation. This approach allows for both spatial (via the 3D printing) and temporal (via the gradient concentration and flow rate) generation of chemical cues in 3D matrices, plus the dynamic regulation of cellular behaviors at a local level. Epithelial growth factor (EGF) is being used, as a proof of concept, to model the migration of cells in the 3D construct toward the chemoattractant. MC7 cells show a different pattern of migration toward the epithelial growth factor gradient in the device when the tumor architecture is different. Combining these 3D bioprinting approaches for in vitro tumor modeling, with microfluidic devices to model microenvironment, provides tools to create complex tumor architecture accompanied by controlled chemical densities, with a high spatiotemporal resolution, beyond what is possible with conventional fabrication technologies.

## Experimental

### Materials

Gelatin from bovine skin (type B) and alginic acid sodium salt, dimethyl sulfoxide (DMSO), calcium chloride, and sodium chloride powder were purchased from Sigma-Aldrich (Canada). Calcium chloride was dissolved in Millipore water at a final concentration of 4%, filtered with 0.2 u filters and stored in a sterile condition for further use. For cell culture studies, MCF7 and MDA-MB-231 cell lines were purchased from ATCC (Cedarlane, Canada). DMEM (Dulbecco’s Modified Eagle’s Medium, without sodium pyruvate, with 4.5 g/l glucose, with L-glutamine), PBS (phosphate buffer saline) 1x, Penicillin-streptomycin, Trypsin/EDTA solution at 0.25% (w./v.) and FBS (fetal bovine serum) were purchased from Wisent Bioproducts (Canada). (3-(4,5-dimethyl-2-thiazol)-2,5-diphenyl-2H-tetrazolium bromide) MTT powder, Triton X-100, BSA (bovine serum albumin), paraformaldehyde, a Live/Dead Cell Viability assay kit (CBA415) and Hoechst 33342 Nuclei Dye, Propidium Iodide (PI) and Calcine were bought from Sigma-Aldrich (Canada). For bioprinting, the mixing syringe (5ml) with the mixing ratio of 4:1, and compatible mixer tips were purchased from Sulzer Inc. (Switzerland). Empty cartridges and nozzles purchased from Cellink (Sweden)

### Preparation of bioink

Sodium alginate and gelatin (type A, 90–110 bloom derived from porcine skin) were purchased from Sigma Aldrich (St. Louis, MO). Bioink formulations were prepared with various ratios of alginate and gelatin. According to the protocol provided by Ouyang et al.,^51^ gelatin and alginate powder mixtures with different ratios were first dissolved in 0.5 % sodium chloride salt solution. Then the solution was stirred vigorously on a hot plate at 85 °C for half an hour. The hydrogels were moved under a biosafety cabinet and treated with UV light for 20 minutes. Afterward, all bioink formations were kept at room temperature (23–24°C) for 2–4 h prior to rheological tests and kept in a fridge (4 °C) for longer storage. All subsequent experiments were conducted at room temperature.

We used two types of cells for our experiments: MCF7 and MDA-MB-231 breast cancer cell lines purchased from ATCC. The cells were passaged once or twice a week and cultured in Dulbecco’s Modified Eagle’s Medium (DMEM) with 10% fetal bovine serum (FBS) and 1% penicillin-streptomycin, called complete DMEM afterward, as recommended by the supplier. Experiments were performed with cells passaged less than ten times. Mixing cells and hydrogel was performed under biosafety cabinet with the dual mixing syringes with the ratio of 4:1 (gel: cell). Cartridges of bioinks containing cells were immediately used for bioprinting under sterile conditions.

### Cell viability and proliferation

Cell viability was evaluated, by MTT and live and dead Assays, on bioprinted structures immediately after printing and after 4, 7 and 11 days. Cell viability properties of constructs were determined using a Live/Dead staining viability kit based on the manufacturer’s instructions. Briefly, each construct was submerged in 1μM Calcine-AM dye solution and incubated for 1 hour at 37 °C. Then the solution aspirates and the constructs were submerged in 2μM propidium iodide solution and kept in the dark at room temperature for 30 minutes. The staining solution was removed, and the constructs were soaked in PBS for 15 minutes twice. A laser scanning confocal microscope (Zeiss LSM 700) was used to image live and dead cells in the constructs at multiple spots. Three constructs (*n* = 3) were used to analyze quantitative cell viability. Based on the live and dead images, the quantitative viabilities of cells were manually counted using ImageJ software (NIH) and calculated based on the number of live cells (green stained cells) over the total number of cells (Live + dead).

Cell proliferation within the 3D network was assayed using MTT assay. 900 μl of DMEM and 100 μl of MTT working solution (0.5 mg mL ^-1^ in PBS) were added to each well of a 12-well plate on each construct and incubated for 3 h at 37°C, 5% CO_2_. At the scheduled timing, the medium was discarded, and cells were washed twice with PBS. Cell membranes and insoluble formazan crystals were dissolved by using 1 ml of dimethyl sulfoxide (DMSO) to each structure for 30 minutes, protected from light. The obtained solution was transferred in a 96-multiwell, and optical density was evaluated at 540 nm using Absorbance Microplate Reader (Biotek Synergy H1, Agilent). Results of cellular viability were expressed as UV-Vis absorbance of each sample compared with the negative control absorbance, which is a 3D construct prototype without cells to exclude the polymer interference in the UV-vis reading.

### Microscopic examinations

All constructs were examined for bright field imaging and fluorescent imaging microscope using Nikon Elipise Ti2 by appropriate filters. A laser scanning confocal microscope (Zeiss LSM 700) was used to observe cell morphology and cell distribution within the 3D printed constructs. At each observation position, a Z-stack scan (400 *μ*m thickness) was implemented with 5 or 10 *μ*m steps at magnifications of ×4.

### Cell Staining

The PKH67 and PKH26 Fluorescent Cell Linker Kits used proprietary membrane labeling technology to stably incorporate a green and red fluorescent dye with long aliphatic tails into lipid regions of the cell membrane.^52^ Diluent C is used as the labeling vehicle to maintain cell viability while maximizing dye solubility and staining efficiency during the labeling step. The cells were centrifuged and suspended in a medium without serum (DMEM only). 4 μL of the PKH67 (green) or PKH67 (red) was added to 1 mL Diluent C and mixed well to disperse. The cell pellets were dispersed in 1 mL of Diluent C. Rapidly, the cell suspension solution and the dye solution were mixed and homogenized by gentle pipetting and incubated at 37 °C for 5 minutes. The staining was stopped by adding 10 mL of complete media (DMEM + 10% FBS and 1% penicillin-streptomycin). Cells were centrifuged and washed with DMEM only two more times to ensure the removal of unbound dye.

### Bioprinting process

CELLINK INCREDIBLE+ (CELLINK AB) was used for the printing of single cell and multiple cell laden constructs. The bioinks were loaded into sterile printing cartridges, and the bioink was extruded through nozzles using extrusion-based print-heads. For sequential printing of two different cells, two separate cartridges have been used. The material flow for each print-head was controlled by individual pressure and speed regulators. The moving speed of the needle was adjusted at 5 mms^-1^. Predefined structures were implemented using Solidwork software and sliced in sli3r software to 10 layers with a rectilinear filling pattern (Figure 1A). The printability of alginate-gelatin hydrogels, using a combination of different hydrogel compositions and printing pressures, was evaluated (Table 1, Supporting Information) using a sterile high precision 22G conical nozzle (0.41 mm inner diameter). All bioink components were sterilised prior to printing by using the UV assembly and soaking in ethanol 70% for 1 hour. The printer, print bed and print-head were kept at room temperature. The bioprinted constructs were subsequently submerged in 4% (w/v) sterile CaCl_2_ for 10 minutes for crosslinking sodium alginate with calcium ions. After 10 minutes, cell-laden constructs were washed three times with PBS, submerged in a growth medium (DMEM) containing 10% FBS and 1% penicillin/streptomycin and incubated (37 °C, 5% CO_2_) for further analysis. The culture medium was exchanged every other day.

**Figure 1:**
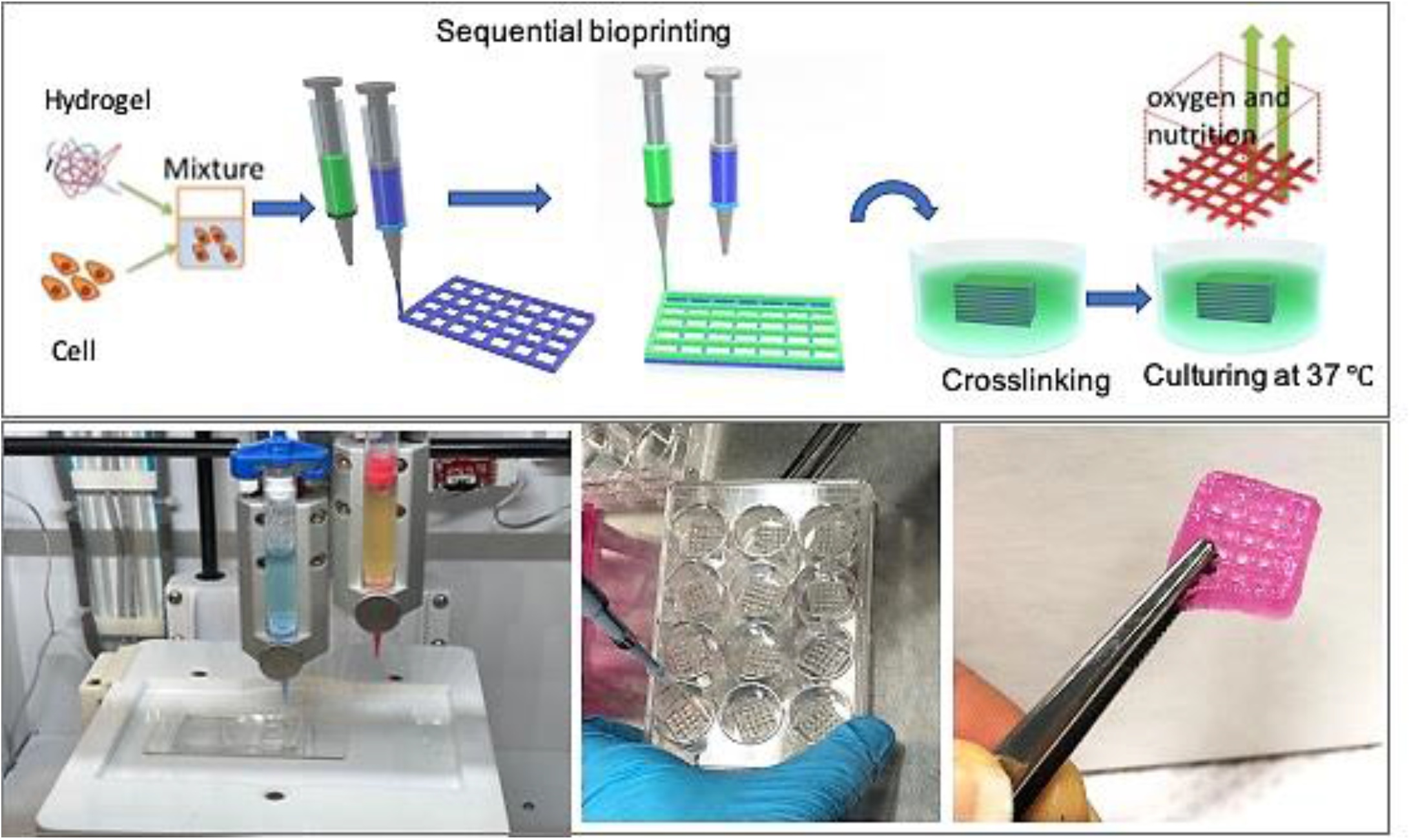
shows (a) the schematic illustration of the biofabrication process, (b) the printer and bioprinted constructs. (Food coloring used for the purpose of illustration)

### Rheological measurements

Rheological measurements of bioink formulations were performed with a Malvern Kinexus Ultra+ rheometer with a 20 mm parallel plate geometry with 1mm gap. To load the hydrogel sample, the hydrogel was placed on the lower plate of the rheometer, and the steel plate geometry was lowered until contact with the surface of the bioink sample. Subsequently, the excess sample around the plate was removed with a spatula. The elastic and viscous moduli of hydrogels were evaluated at 25 °C and 15 °C at a strain of 0.1 %. Viscosities of all samples were measured at 25 °C at a shear rate of 1 to 100 s^-1^. The shear elastic modulus *G′*, shear loss modulus *G″*, and loss tangent tan(δ) were measured for each hydrogel bioink using a shear strain sweep test ranging from 0.02% to 1.0% at an oscillation frequency of 40 Hz. All rheological measurements were conducted in triplicate.

### Microfluidic chip design and manufacturing

#### Design

A tree-like gradient generator is designed that can be integrated with the 3D bioprinting for the tumor-on-chip model. The device contains two inlets for the injection of the culture media as well as the reagent of interest. A linear gradient of the reagent will be produced across a chamber while the waste materials leave the device through six outlets next to this chamber. An algorithm was utilized to find the length of branches in each step of mixing channels, assuming the width of channels and their height were 400 μm and 50 μm respectively. The whole model is 21 mm in width and 45 mm in length.

#### Fabrication

A standard lithography procedure was followed to fabricate a microfluidic gradient generator. Briefly, a 50μm thick layer of SU8-2050 negative photoresist was spun on a cleaned Si wafer and then was patterned and developed by rinsing in SU8 etchant to form the mold for the microchannels. Next, a 10:1 ratio of PDMS was spread onto the mold and was left for 12 h at 65 ⁰C to completely polymerize. The cast chip in PDMS was peeled off, and holes were punched in the location of the in/outlets. The chamber area was also cut out using a blade to open some space for the tumor printing. Then, the PDMS layer was bonded to a glass slide using oxygen plasma in plasma cleaner. The as-prepared microfluidic chip was exposed to UV light for disinfection before bioprinting tumors directly in the chamber or transferred to the chamber. Another glass slide was activated using oxygen plasma and sealed the top of the chamber. A flow of cell culture media using a syringe pump was immediately introduced through the inlets to provide cell viability. The source in one of the inlets was then switched with the reagent of interest to form the gradient in the chamber. The device was kept inside an incubator at 37⁰C and 5% CO_2_ during the whole experiment and was periodically examined.

### Tumor-on-chip modeling

Three models were built in this study, including a 2D model, a sequential 3D bioprinting model and a co-cultured 3D bioprinting, all in on the microfluidic device. The 2D model was prepared simply by culturing MCF7 and MDA-MB-231 cells in 12-well plates to 80% confluency unless otherwise stated. For co-culture experiments, the cells were labelled with different florescent markers before culturing to distinct visibility by fluorescent microscope during the experiment. For sequential bioprinting, a G-code has been prepared for the bioprinter to allow switching between two printheads in alternative layers in which each layer was printed with one of the bioinks including type A or type B cells. For the mixed cell bioprinted constructs, two different cells were prepared in labelled with green and red markers to be distinguishable. The hydrogel was mixed with cells with the desired ratio of two cells and desired ratio of gel to cells. One cartridge of mixed cells was prepared and printed in a layer-by-layer fashion. For the microfluidic chip experiments, the constructs can be printed directly in the microfluidic chamber or transferred to the chamber later, based on the designed test.

## Results

### Printability and mechanical properties of bioink

In the preliminary study, various hydrogels with alginate 1-10% and gelatin 4-10% were prepared and printed at room temperature using INKREDIBLE bioprinter (Cellink, Sweden). The small nozzle diameter results in higher resolution, but increases cellular damage because of high extrusion pressure. In the other hand, previous studies reported that low shape fidelity obtained with higher nozzle diameter with low pressure printing.^16^ However it provides a better condition for cell viability. A quantitative relationship between nozzle size and dispensing pressure on cell viability has been found.^41^ Here, each hydrogel was printed with two nozzle diameters, 25 G (Red, 250 µm inner diameter) 22G (Blue, 410 µm inner diameter). Figure 1 shows the schematic illustration of the biofabrication process and printed structures. We continued bioprinting with 22G nozzle that needs less extrusion pressure and was found favorable for cell viability after printing. After making bioink by mixing hydrogel and the cells, the bioprinting procedure was applied sequentially by two print heads containing cells A and B separately. Then the printed construct was submerged in CaCl_2_ for crosslinking for 10 minutes. After washing by PBS, each construct was immersed in cell media (DMEM, with 10% PBS, 1% Pen-Stripe) and incubated in 37 °C. Table 1 (Supporting Information) summarizes the applied pressure, filament condition and quality of the printed structure based on nozzle size and hydrogel composition.

The rheological properties of hydrogel samples for four combinations of A1G4, A1G8, A4G4 and A8G4, (which A1G4 represents Alginate 1%, Gelatin 4%, etc) with the best printability have been studied. Figure 2 shows the rheology characterization of hydrogel mixtures. Figure 2A, shows the storage modulus (G’) and loss modulus (G”) obtained at room temperature by frequency oscillation measurement. The viscoelasticity data for all samples show a higher storage modulus than loss modulus, which indicates the gel-like property of the material. Both storage and loss modulus were found to increase as frequency increased.

**Figure 2:**
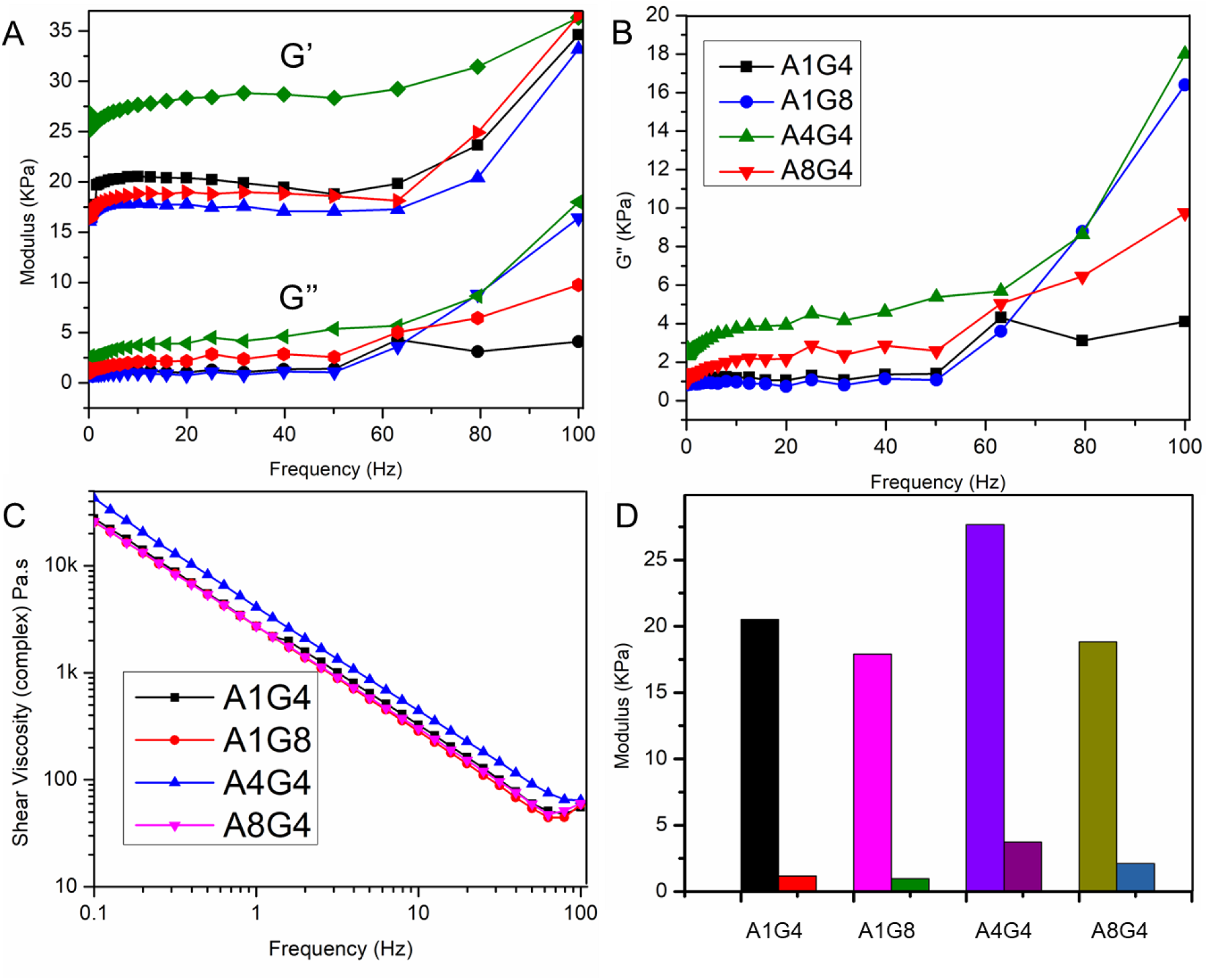
Rheology characterization of hydrogel mixtures with A1G4, A4G4, A8G4 and A1G8 (1, 4, 8, and 1 % alginate and 4, 4, 4, and 8% gelatin respectively). (A) Storage and loss modulus of A1G4, A1G8, A4G4 and A8G4 under different oscillatory frequencies. (B) Tan δ vs frequency of the hydrogel mixtures. Tan δ value were found <1 for all hydrogel mixtures. (C) Viscosity of hydrogels at different shear rate. Viscosity of all hydrogels decreased with increasing shear rate. (D) Histogram of modulus for different hydrogel mixtures. A4G4 shows the highest storage modulus. (D) loss modulus, G”

Tan δ values were calculated and are shown in Figure 2B versus frequency for four hydrogels. Tan δ values were found <1, which suggested that all the hydrogel mixtures are more gel like mixture than liquid like. The strength of hydrogel was increased as alginate concentration increased up to alginate 4% and dropped when alginate concentration is 8%.

The viscosity of all hydrogels mixtures, shown in Figure 2C, decreased with increasing shear rate and the shear-thinning behavior was observed for all four samples. The viscosity curves of all the hydrogel mixtures showed a similar pattern which suggests that all the hydrogels had shear thinning behaviors. The loss and storage modulus at 40 Hz are illustrated in Figure 2D. A4G4 shows the highest storage (elastic) modulus. It has also been observed that 4% gelatin contributes a profound viscoelastic property when mixed with 4% alginate, which resulted in consistent extrusion through the nozzle.

### 1.1. Cell viability

To visualize the distribution of cells in printed construct, cell membranes were pre-stained with a green fluorescent membrane marker (PKH67) before printing and imaged immediately after printing with a confocal microscope. Figure 3A shows that cells are distributed homogeneously in the construct. The viability of different cells after printing was determined using a live-dead staining assay. The live/dead staining reagents were added to each structure immediately after bioprinting, as described in the experimental section. The status of cell survival was defined as the percentage ratio of the live cells over the total cells, where calcium-acetoxymethyl green fluorescence stained live cells and propidium iodide red fluorescence stained dead cells. Cell viability is an essential factor for the successful fabrication of cell-printed constructs. The live-dead assay results (Figure 3B) show the percentage of viable MDA-MB-231 cells immediately after printing 94.16%, 96.94%, 75.96%, and 84.7% for hydrogels A1G4, A4G4, A8G4 and, A1G8 respectively. The representing image related to A4G4 with green shows live cells and red indicates dead cells presented in Figure 3C. The 3D z-stack is shown in Figure 3D, confirming the homogeneity of live cells in the construct.

**Figure 3:**
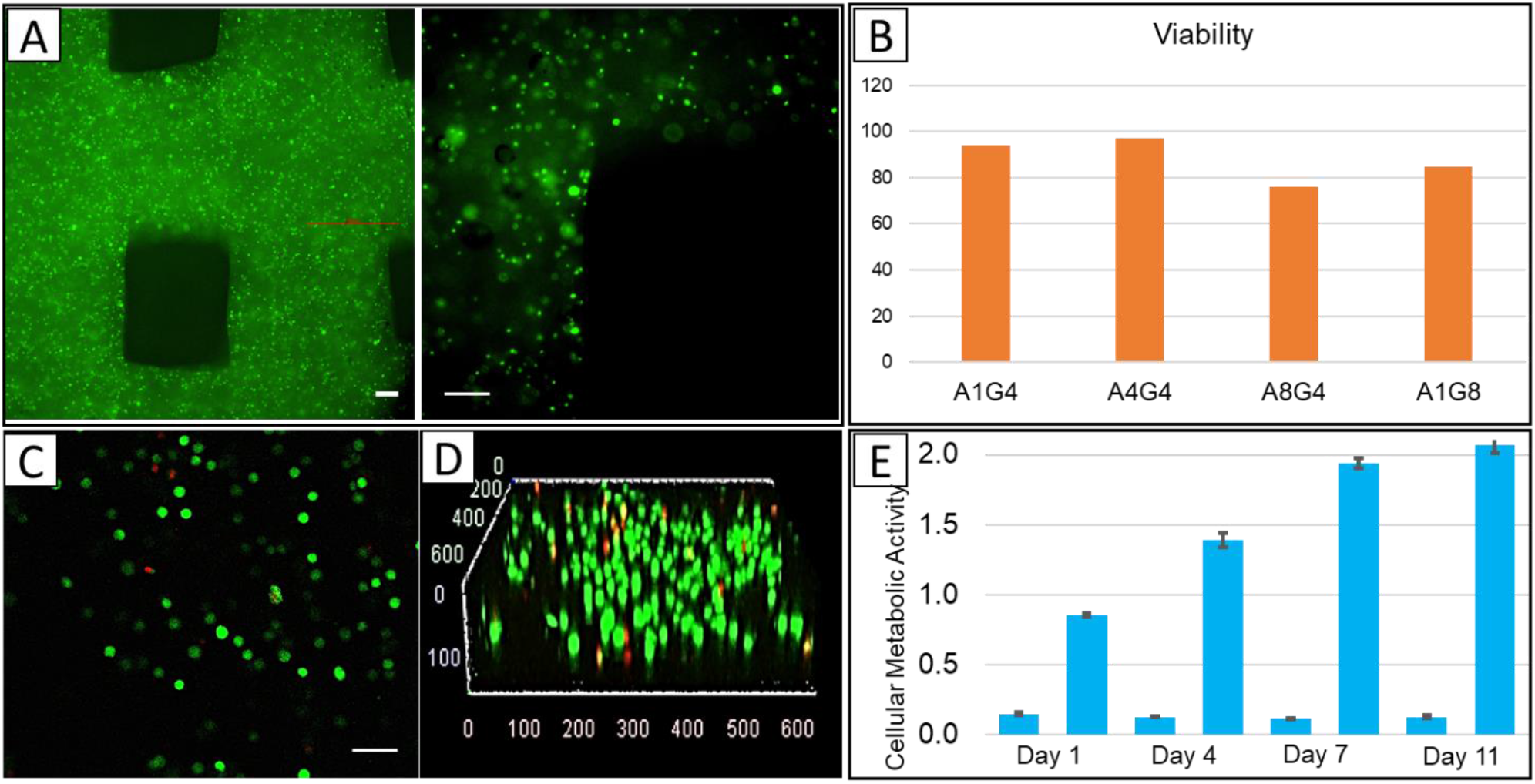
The uniform distribution and viability of bioprinted MDA-MD-231 cells. (A), Concentration and distribution of printed cells in the construct, Scale bars, 50 μm (B) Histogram shows the viability of the cells in the hydrogel mixtures immediately after printing. (Status of cell survival by the percentage ratio of live/dead cells) Live-dead staining of four different hydrogel mixtures with MDA-MB-231 cells after bioprinting. (C) The representing image related to A4G4 with the green shows live cells and red for dead cells, Scale bar, 50 μm (D) 3D confocal representative image for A4G4 hydrogel. (E) Results of MTT assay on bioprinted constructs with A4G4 hydrogel, immediately after printing (day 1), after 4 days, after 7 days and after 11 days. Data are expressed as the average ± standard deviation of three replicates of samples and three replicates of negative controls (small bars).

3-(4,5-Dimethylthiazol-2-yl)-2,5-diphenyltetrazolium bromide (MTT) assay was conducted on three replicates of 3D bioprinted constructs immediately after printing and after incubation at 37°C, 5% CO_2_ in DMEM with 10% FBS and 1% Penicillin-streptavidin for 4, 7 and 11 days. Results of MTT test are reported in Figure 3E as absorbance data detected at 570 nm. The data refer to three replicates of 3D bioprinted constructs and three replicates of negative control constructs (small bars) that are 3D printed structures without cells. A clearly good cell distribution into construct can be observed with absorbance values were ∼ 0.86 ± 0.02 immediately after printing. An increment of absorbance was measured after incubation for 4 days (1.39 ± 0.05) and continued to reach 1.94 ± 0.04 on day 7. The rate of viability, compared to day 1, increased over the week but on the day 11, the absorbance was 2.07± 0.06, which shows a drop in the rate of cell proliferation. It seems that between day 7 to day 11 the constructs reach the maximum capacity for hosting cells and some cells started to die.

### 1.2. 3D Bioprinted co-cultures

MCF7 cancer cells (a breast adenocarcinoma cell line) and MDA-MB-231 (a more aggressive triple negative breast cancer cell line with mesenchymal characteristics) are pre-stained with green and red markers before printing, respectively. In the first printing procedure, two bioinks were prepared separately from each type of cell and printed sequentially. The first layer of the printed structure consisted of MDA-MB-231 cells (stained red), and the second layer of MCF7 cells (stained green) lay on top of the first layer. In the second printing experiment, one bioink was prepared with both pre-stained cells mixture in the hydrogel and bioprinting was conducted layer-by-layer same as before. Figures 4A and 4B show the confocal image of the sequential bioprinted cells and bioprinted structure consisting of a mixture of both cells, respectively. In a separate experiment, the mixture of MCF7 and MDA-MB-231 cells was co-cultured in a 2D chamber. It can be seen that the cell population and the cell ratio are different at the edges of the chamber compared to the middle of the chamber, while in 3D bioprinted structure, the cells are located homogeneously. (Figure 4C, 4D and 4E) It should be noted that single cell cultures of MCF7 and MDA-MB-231 cells were prepared with and without cell staining to observe the effect of staining on cell growth. The co-culture with a mixture of both cells was prepared in both conditions. The cell growth was observed under a microscope after 3 days and 5 days. No noticeable differences were observed between cultures from cells with staining and without staining, shows the staining procedure does not have an adverse effect on cell growth. (data not shown here)

**Figure 4:**
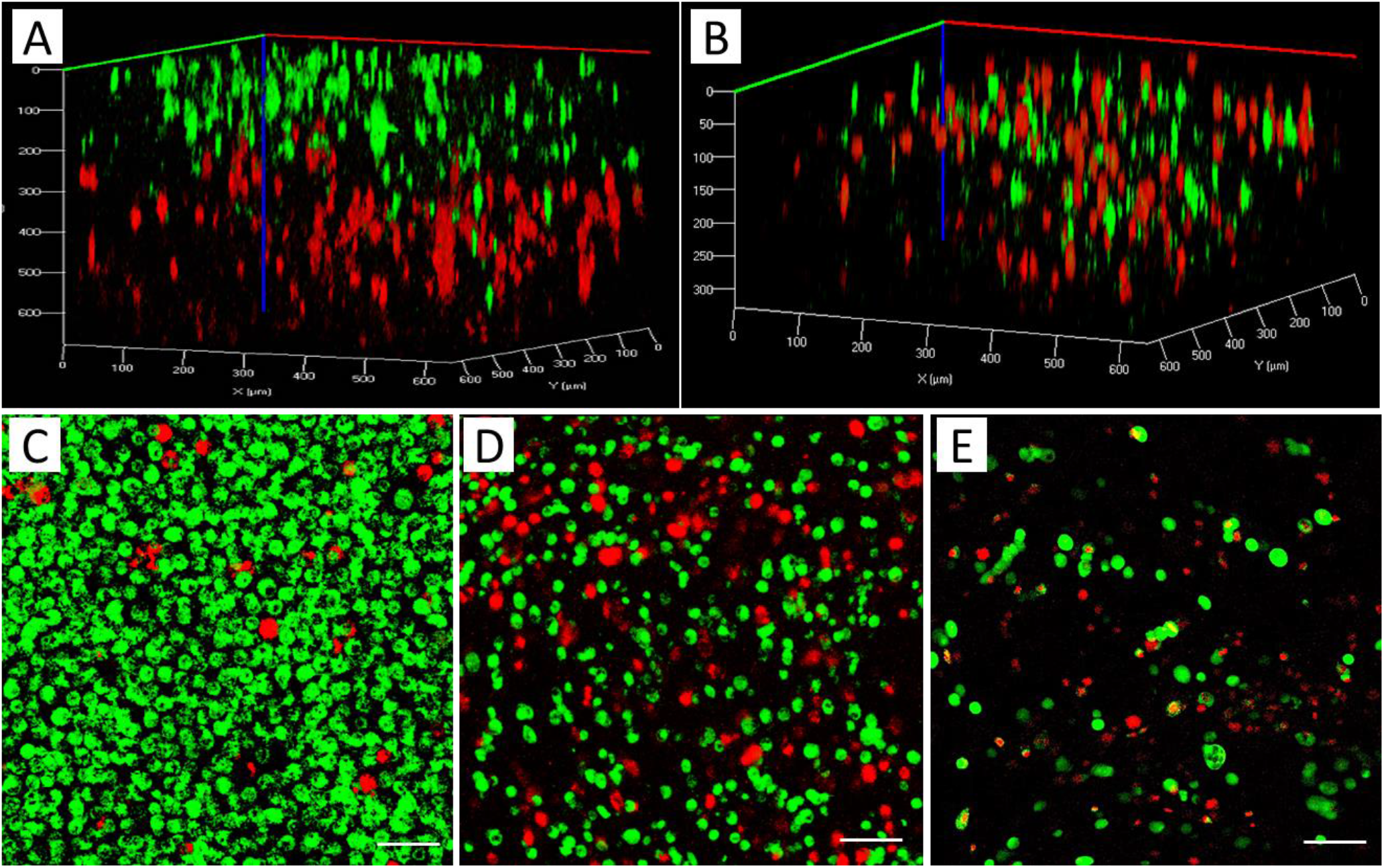
3D rendered confocal images of MCF7 (stained green) and MDA-MB-231 cells (stained red): A) Sequential printing of MDA-MB-231 and MCF7, B) bioprinting of a construct using a bioink with a mixture of MCF7 and MDA-MB-231 cells; 2D culture of both cells mixture with the ratio of 2:1 (MCF7, green:MDA-MB-231, red): C) Edge of the culture chamber, D) Middle of the chamber, E) 3D bioprinting of mixture cells with the ratio of 2:1 (green:red). Scale bars, 50 µm.

In order to investigate cell growth and aggregation in bioprinted co-cultures, 3D construct consisting of each cell type was separately printed and compared with the construct consisting of both cells’ mixture in the bioink. A confocal microscope was used to observe the morphology of MCF7 and MDA-MB-231 cells over a 10-day experimental period in the tumor-like constructs. (Figure 1S, Supporting Information). Confocal imaging of co-culture 3D bioprinted constructs shows that at day 1, both MCF7 (red) and MDA-MB-231 (green) cells are present in each layer randomly as they printed. After 3 days, both cell lines can be observed, but it seems the cells are in different layers and migrated toward the same phenotype to make cell clusters. It seems the migration rate of MCF7 inside the hydrogel is a way that most of the cultures are either MDA-MB-231 or a mixture of MDA-MCf7. Few clusters of MCF7 cells are also observed. The cell migration inside the hydrogel is consistent with previous reports investigating the pore size of alginate-gelatin hydrogels. ^46,47^

### 1.3. Cell behaviour in response to chemoattractant

Here, we used epithelial growth factor (EGF), a chemoattractant material, to observe cell migration in co-cultures of the microfluidic device. The extracellular biomolecular gradients are dynamically generated within 3D hydrogel matrices via a linear gradient designed to mimic chemical environments in tumor tissues and direct cell migration. An algorithm was utilized to find the length of branches in each step of mixing channels, assuming the width of channels and their height were 400 μm and 50 μm, respectively. The whole model is 21 mm in width and 45 mm in length. Figure 5A shows the COMSOL simulation of this device in which Navier-stokes and Convection-Diffusion equations are solved using the finite element method. In this figure, red and blue colors represent higher and lower concentrations in [mol/mm^3^], respectively. The diffusion coefficient was considered to be 10^9^ m^2^/s which is a typical value in aqueous solutions. Pure water as the solvent and 1 [mol/mm^3^] aqueous solution was supposed to enter the chip through inlets 1 and 2, respectively.

**Figure 5:**
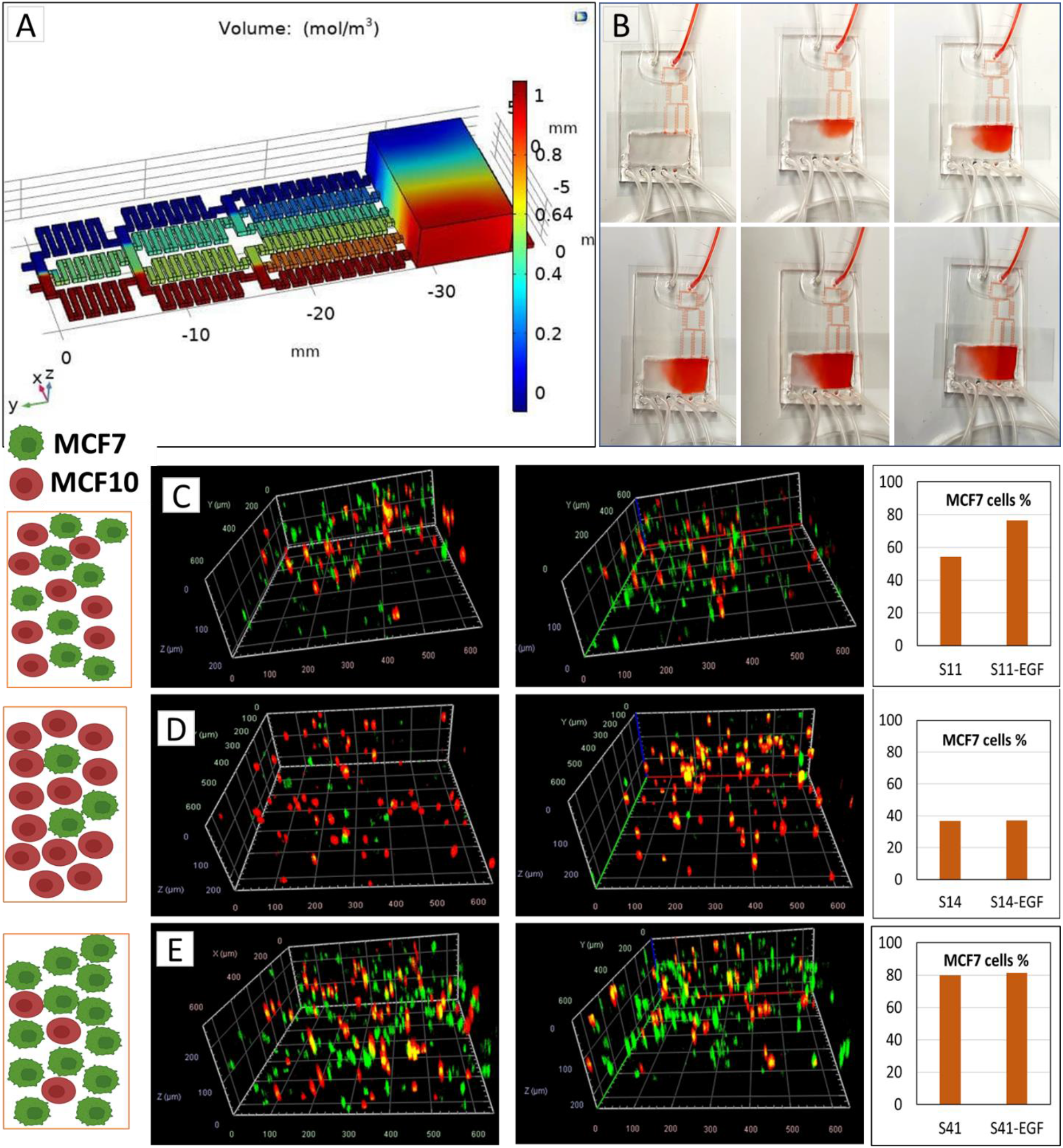
(A) COMSOL simulation of the device with red and blue represent higher and lower concentrations, (B) gradient formation in the 5 min intervals. The migration of MCF7 (green) and MCF10 (red) toward EGF in the microfluidic chamber; imaged on the higher concentrated of the chamber before and after EGF gradient flow: MCF7/MCF10 ratio 1:1 (S11) (C), MCF7/MCF10 ratio 1:4 (S14) (D) and MCF7/MCF10 ratio 4:1 (S41) (E). The respective percentage of MCF7 cells in before and after EGF exposure are shown for each panel.

To confirm the chemical gradient, the cell culture medium and FBS were used as the source on the inlets to produce diffusion visual confirmation. Figure 5B shows the picture of the chamber with 5 minutes intervals applying a flow rate of 20 μl/min. The gradient has been produced and obtained for at least 5 hours without the flow. Bioprinted constructs, including MCF7 (labelled green) and MCF10 (labelled red) were placed in the gradient chamber. The flow rate of 5 μl/min was used with the cell culture media as the source for outlet 1 and cell culture media with 100 nM EGF as the source of outlet 2. The gradient was introduced to the cells for 8 hours while the whole device was kept in 37 °C with 5% CO_2_. Figure 5C, 5D and 5E shows different tumor architectures of MCF7 and MCF10 cells bio-printed with ratios of MCF7/MCF10: 1/1 (S11), 1/4 (S14) and 4/1 (S41). The left panels show the MCF7 and MCF10 distribution in the tumor before applying EGF and the right panels shows their distribution after applying EGF. For each panel the total percentage of MCF7 cells before and after EGF exposure has been calculated and shown in the plots. The highest migration tendency observed when the ratio of cancer cells to the surrounding healthy cells is 1:1. (Figure 5C)

## Discussion

Tumor heterogeneity has been observed among patients. Breast tumors are heterogeneous and consist of many different cell types that create the TME. Tumor development and the microenvironment cells have a two-way influence on each other by secreting cytokines, growth factors, etc. Besides the cancer cell stromal cells, individual tumors show heterogeneity, which was first observed about four decades ago in murine models^23^ and it is named intratumor heterogeneity. Within individual cancers, there are distinct cancer cell subpopulations with variable metastatic ability due to their different representations of tumorigenicity, signaling pathways, migration and response to anticancer drugs.^24,25^ As a result, an effective model must not only provide a microenvironment for 3D growth of cells but also allow cell–cell interactions to study cancer progression. Any tumor-on-chip model should cover the tumor heterogeneity to be accurate.

Various 3D tissue models is created before to study two of the most important factors in cancer mortality, angiogenesis and metastasis.^26^ In many metastases cases, death results from distant metastases to other organs and tissues.^27,28^ Angiogenesis modeling is also a critical factor in tumor development and metastasis, allowing investigation of vascular endothelial cells’ interactions with cancer cells, including the co-culture of tumor cells with endothelial cells, explanted vessels or seeded cells surrounded by tumor cell-laden hydrogels.^29–31^ Here, our goal is to establish a platform using 3D bioprinting for modeling the phenotypic heterogeneity, which is the result of different cell localization in a tumor using bio-printed tumors.

The material’s intrinsic properties (including modulus, yield stress and viscosity) is the most important factor for its printability and some other external conditions are also important such as applied pressure, nozzle geometry.^32^ The key component of a successful tumor 3D printing is developing an appropriate bioink. Hydrogels, with about 99% water content, are primary candidates for bioprinting since their high amount of water favors the entrapment of biological entities and increases hydrogel biocompatibility.^33^ Moreover, they have significant impacts on cell proliferation, migration, aggregation, and activities that are strongly dependent to hydrogels’ physical properties.^34^ To date, various types of synthetic and natural polymers have been used for hydrogel preparations with different biocompatibilities and mechanical properties.^34^ Polymers with low cytotoxicity and structural similarity to the ECM are good candidates for making bioinks. The primary requirements for a bioprinting material are a) cell viability, b) printability (shear-thinning, compatible printing pressure, continuous extrusion without air bubbles), c) structural stability (stability of a printed structure before crosslinking and after crosslinking), and d) rapid and non-toxic crosslinking.^35^

The ideal bioink for bioprinting with extrusion methods should have relatively high viscosity, show strong shear-thinning behavior and rapidly cross link after printing. Therefore, natural polymers like alginate, gelatin, chitosan and hyaluronic acid hydrogels have attracted more attention due to their high compatibility to cell environments and good control over the quantity of ECM proteins and growth factors.^35^ A widely investigated hydrogel is alginate, a natural polysaccharide derived from algae which is unique because it undergoes a sol-gel transition in the presence of divalent cations such as calcium. It also has optimal cell encapsulation properties as the slow gelation process at room temperature.^36,37^ However, alginate shows poor mechanical properties. Gelatin is a highly elastic material rather than viscous and is considered to be a stiff solid bioink at room temperature^8^, which is not ideal for extrusion bioprinting methods. On the other hand, if gelatin is mixed with alginate hydrogels to form a liquid phase gel, an optimum condition for bioprinting by extrusion can be achieved. Previous reports studied combinations of gelatin and alginate compatible with extrusion bioprinters.^35,38^ They have also shown that gelatin provides elastic characteristics in the hydrogel, improves cell adhesion, contributes to the viscosity of the hydrogel, and adds stiffness.^39,40^

A high shear stress can cause damage to the cell membrane. Therefore, the shear thinning ability is important to reduce the shear stress imparted on the cells during the extrusion process. A bioink to be considered printable should provide the opportunity for the fabrication of multi-layer porous structure without collapsing or sagging after printing. For our ink to be printable, it should exhibit sufficient yield stress to prevent its collapse, has smooth extrusion out of the nozzle such that no corrugation appears and should not show “peaks and valleys” along the extrudate or breaks within one filament.

We developed various hydrogels with alginate 1-10% and gelatin 4-10% and printed them at room temperature. By increasing the concentration of alginate from 1% to 8% while keeping the gelatin concentration about 4%, the filament quality improves, which is consistent with the report of Giuseppe ^42^ that increasing the concentration of alginate and gelatin resulted in more accurate printing. However, increasing the concentration of gelatin by more than 4% reduces the accuracy of the printing. It also needs higher extrusion pressure due to nozzle jamming. On the other hand, the synergetic properties of alginate-gelatin made Alginate 1%- Gelatin 8% still a good candidate for accurate printing.

Several criteria for bioink mechanical properties can be considered, but the most important one is rheology.^8^ Rheology is the measurement of the deformation of a material caused by force acting on it and the vast majority of rheological characterizations of bioinks have focused on hydrogel viscosity. The rheological properties of hydrogel samples for four combinations of A1G4, A1G8, A4G4 and A8G4, (which A1G4 represents Alginate 1%, Gelatin 4%, etc) with the best printability have been studied. Hydrogels with higher viscosity, experience high shear forces to extrude through the nozzle during printing. Shear thinning behavior enables highly viscous hydrogels printable with structure accuracy.^43^ As it can be seen in Figure 2C, viscosity of all hydrogels mixtures, decreased with increasing shear rate and the shear-thinning behavior was observed for all four samples. The viscosity curves of all the hydrogel mixtures showed a similar pattern which suggests that all the hydrogels had shear thinning behaviors. Based on the loss and storage modulus of hydrogels illustrated in Figure 2D, A4G4 shows the highest storage (elastic) modulus.

The composition and mechanical properties of the hydrogel play an important role in cell viability. Based on the live-dead assay results that previously shown in Figure 3B, the percentage of viable MDA-MB-231 cells immediately after printing 94.16%, 96.94%, 75.96%, and 84.7% for hydrogels A1G4, A4G4, A8G4 and, A1G8 respectively. A droplet of the A1G8 hydrogel, as a representative of not printed gel, was placed in CaCl_2_ for crosslinking and underwent to live-dead assay process. The cell viability for A1G8 before printing was also obtained as 93.4% that is slightly higher than bioprinted one (84.7%), which is in accordance with the extrusion procedure affecting cell viability. A1G4 and A4G4 show the best cell viability results, however, A1G4 did not have a good shape fidelity in the hydrogel structure. Therefore, based on printability and viability results, A4G4 was chosen as the optimized hydrogel to be used for the co-culture experiment. Excellent viability obtained from a mixture of 4% alginate and 4% gelatin can be attributed to the soft nature of gel and cell adhesion that gelatin introduced in the hydrogel. Previous reports confirmed that when alginate concentrations is decreasing in hydrogel compositions and gelatin concentration is increasing, the gels are mechanically soft and contain a greater number of cell-adhesion moieties.^44^ Our results are consistent with other reports that highly viscous hydrogels result in lower cell viability due to high extrusion pressure, which imparts higher shear stress to cells.^40,45^

The MTT assay that conducted on the samples immediately after printing, and after 4, 7, and 11 days indicates the rate of viability in the 3d bioprinted constructs.(Figure 3E) The rate of viability, compared to day 1, increased over the week, which confirms the accessibility of the cells to oxygen and nutrients. The hydrogel constructs had enough porosity and interconnected pore structure, which ensures nutrient diffusion within the construct and provides an appropriate environment for living cells. Moreover, this interconnected porous structure also allows waste product diffusion from the construct. On day 11, the absorbance was 2.07± 0.06, which shows a drop in the rate of cell proliferation but still very good cell viability. It seems that between day 7 to day 11 the constructs reach the maximum capacity for hosting cells and some cells started to die. These findings confirmed good biocompatibility of 3D bioprinted hydrogel constructs and showed their suitability for preserving cells viable for a longer time.

For the fabrication of co-cultures by bioprinting two cell lines, MCF7 cancer cells and MDA-MB-231 are pre-stained with green and red markers before printing, respectively. The printing procedure conducted in two ways to produce two different architectures, one including two separate layers of cells and the other one randomly mixture of two cells. (Shown in Figures 4A and 4B). It can be observed that good control of positioning cells in the construct has been achieved also cells are located homogeneously in the structure compared to the co-culture produced in 2D culture flask. (Figure 4C, 4D and 4E).

Tumour-on-chip models are shown more efficient models in cancer research compared to the other, because of capability to mimic tissue-tissue interactions and tumour vascularization.^48^ Developing these models is an advanced step of bioprinting and provides high-throughput testing, better microfluidic simulations, therefore better in vitro and in vivo correlations are seen. As stated in the introduction, the metastatic spreading of a tumor begins with the acquisition of an aggressive phenotype by a subset of cells which allows them to detach from the primary tumor and migrate toward a secondary organ. Generally, breast cancer metastatic activities increase with cell-matrix interactions, mechanical stimulation and cancer cell interaction with fibroblasts and immune cells.^49,50^ Therefore, for the development of metastasis activity in the designed 3D tumor, the development of a heterogeneous multicellular tumor should be considered.

Several subtypes of breast cancers can be defined, with different tendencies to form metastases into bone.^26^ In this study, we used epithelial growth factor (EGF), a chemoattractant material and observed the cell migration in co-cultures of the microfluidic device. Our 3D microfluidic models have been developed to recapitulate TME heterogeneity by introducing chemical gradient, and we introduce cancer modeling by combination microfluidic device and 3D bio-printing for the first time. To validate the model, breast cancer cell lines (MCF7) and non-tumorigenic mammary epithelial cells (MCF10) were embedded separately within the tumor model, all of which maintained high viability throughout the experiments. (data not shown here) Then, three different tumor architectures of MCF7 and MCF10 cells bio-printed with the following ratios MCF7/MCF10: 1/1 (S11), 1/4 (S14) and 4/1 (S41) and samples were placed in the microfluidic device chamber. The chamber was sealed, and the chemical gradient of EGF was produced for 8 hours. Samples were imaged before and after applying the flow and making the EGF gradient. We compared the ratio of MCF7 cells to MCF10 cells on the top part of the scaffold which was exposed to the highest EGF concentration. MCF7 exhibited migratory behavior toward EGF and many cells were observed to move to the top of the chamber where the EGF concentration was higher. (Figure 5C, 5D and 5E, the second panels) However, MCF10 surrounding cells showed little net migration. The migratory behavior was different when MCF10 cells were present in the co-culture, and the ratio of cancer cells to healthy cells was different. We observed the highest migration tendency when the ratio of cancer cells to the surrounding healthy cells is 1:1, shown in Figure 5C. When the surrounding MCF10 is four times bigger than cancer cells, the migration is negligible, and when the cancer cells are four times more populated than the surrounding cells, the migration rate is decreased. That can be related to the competition of MCF7 cells toward EGF and the accumulation capacity of the scaffold.

### Conclusion and future outlook

We successfully implicated a tumor-on-chip model based on 3D bioprinting of two different types of breast cancer cells and non-tumorigenic mammary epithelial cells. It is crucial to understand the molecular and cellular mechanisms of tumor heterogeneity for development of treatment resistance. Since breast cancer is heterogeneous in both cell lines and in breast tumors, investigation of tumor heterogeneity has important implications for the diagnosis, therapeutic treatment and chemoresistance behaviour. Only in one breast tumor, different types of cancer cell populations are exist which complicates the correct identification of disease subtypes. Furthermore, many studies show that tumor cell-cell interaction and tumor-TME interactions have significant influence on tumor progression and invasion. Our heterogeneous tumor-on-chip model can have huge impact in cancer treatment. In this work, we developed 3D bioprinting methods for building in vitro models of the TME made of co-cultured cells with controlled distribution and architecture in a hydrogel matrix. The tumor was represented by cancer cell-laden co-culture hydrogel construct, whereas its microenvironment modeled in a microfluidic chip capable of producing a chemical gradient. We investigated a composite hydrogel as a bioink, comprised of alginate and gelatin, and an optimized bioink was used for bioprinting the co-cultured constructs. The morphology and distribution of two cancer cell types, MCF7 and MDA-MB-231 were investigated by sequential bioprinting and mixed bioink bioprinting to show the accuracy of cell locations after printing. Epithelial growth factor (EGF) is used as a proof of concept to study cell migration in a different 3D compositional and structural format, including mixed co-culture systems of non-tumorigenic mammary epithelial cells, MCF10 and malignant MCF7 cancer cells.

Better understanding of the relationship between intratumor heterogeneity and tumor response to therapy will open a new approaches for development of new drugs and to use already approved drugs in new treatment schemes or combinations for more effective personalized therapy. Our tumor-on-chip model can facilitate an understanding of cell behaviour in heterogenous tumor and its microenvironment.

## Abbreviations

TME: tumor microenvironment
2D: Two-dimensional
3D: Three-dimensional
ECM: extracellular matrix
EGF: Epithelial growth factor
DMSO: dimethyl sulfoxide
DMEM: Dulbecco’s Modified Eagle’s Medium
MTT: 3-(4,5-dimethyl-2-thiazol)-2,5-diphenyl-2H-tetrazolium bromide
FBS: fetal bovine serum

## Acknowledgment

Not applicable.

## Authors’ contributions

NM designed the experiments, analyzed data, and wrote the manuscript. SAH designed the microfluidic device and relevant simulations. ABD and DM helped in cell culture experiments. AG and MK supervised the research. All authors read and approved the final manuscript.

## Availability of data and materials

All data and materials are available upon request.

## Declarations

### Ethics approval and consent to participate

Not applicable.

### Consent for publication

All authors read the manuscript and agreed to publish it in Biomaterials Research.

---

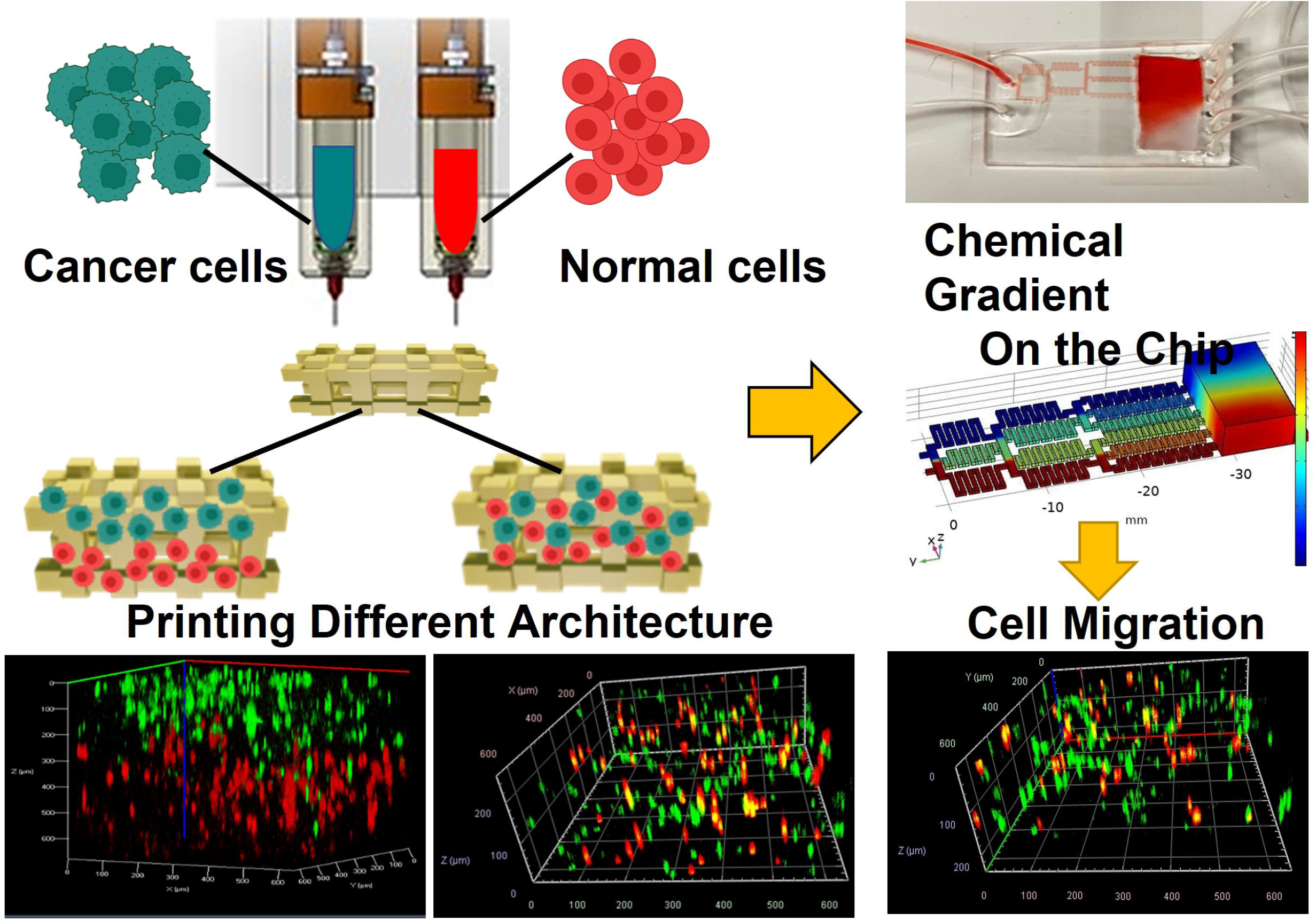

