## Supporting Information, Table 1 for "Controlled tumor heterogeneity in a co-culture system by 3D bio-printed tumor-on-chip model"

| **Bioink** | **Pressure, Gauges (22G Nozzle)** | **Filament** | **Printability** |
|---|---|---|---|
| **Alg 1%/ Gel 4% (A1G4)** | 20-30 KPa | Has nods with red nozzle, spreads with blue nozzle | Could not hold the structure, especially with blue nozzle |
| Alg 1%/ Gel 5% (A1G5) | 20-30 KPa | Has nods with red nozzle, spreads with blue nozzle | Could not hold the structure, especially with blue nozzle |
| **Alg 4%/ Gel 4% (A4G4)** | 60-70 KPa | smooth | Accurate printing |
| **Alg 8%/ Gel 4% (A8G4)** | 100-110 KPa | Very sharp and smooth | Accurate printing |
| Alg 4%/ Gel 8% (A4G8) | 130-140 KPa | Has nods | Lack of printing accuracy, too dense and difficult to print |
| **Alg 1%/ Gel 8% (A1G8)** | 70-80 KPa | smooth | Accurate printing |
| Alg 1%/ Gel 10% (A1G10) | 100-110 KPa | Has nods | Lack of printing accuracy |

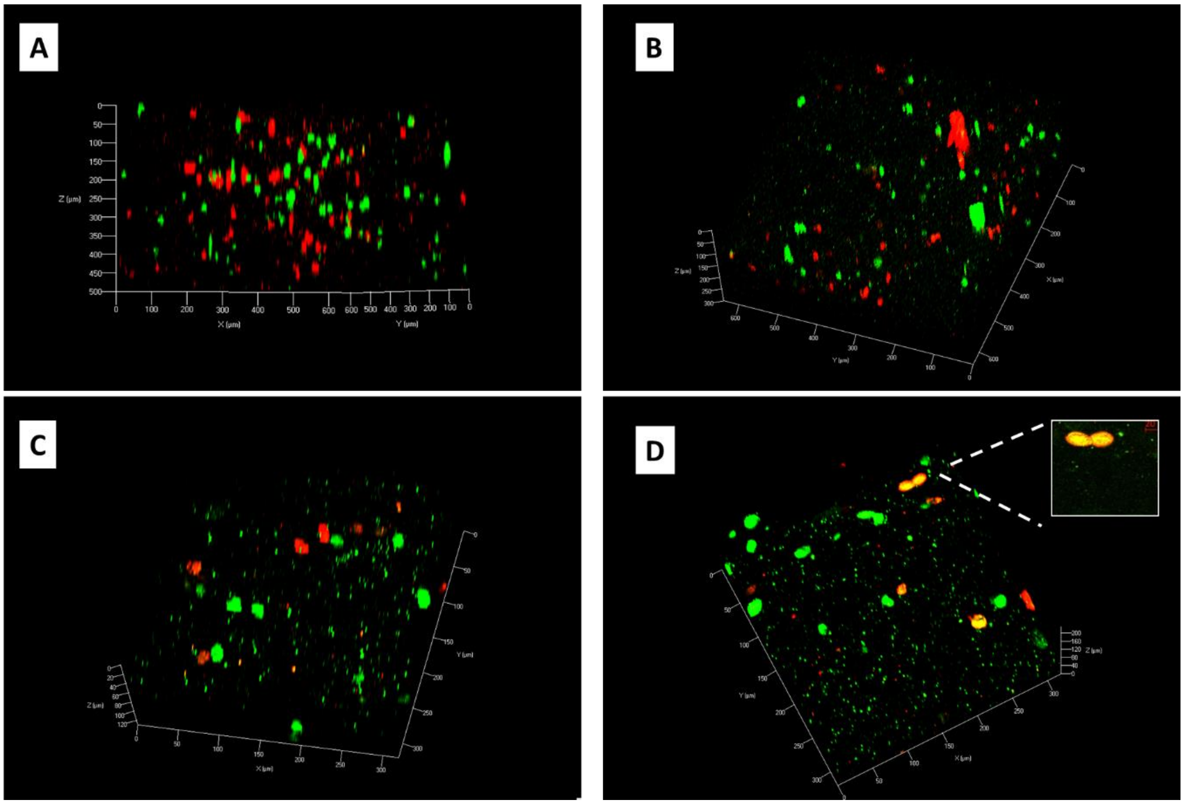


**Figure 1S**: Confocal image of the co-culture construct at (A) day 1, (B) day 10. (C) and (D) are the higher magnification of two different layer of the construct at day 10. MDA-MB-231 cells marked as green and MCF marked as red.
